# Go Get Data (GGD): simple, reproducible access to scientific data

**DOI:** 10.1101/2020.09.10.291377

**Authors:** Michael J. Cormier, Jonathan R. Belyeu, Brent S. Pedersen, Joseph Brown, Johannes Koster, Aaron R. Quinlan

**Affiliations:** Department of Human Genetics, University of Utah, Salt Lake City, UT, USA; Utah Center for Genetic Discovery, University of Utah, Salt Lake City, UT, USA; Institute of Human Genetics, University of Duisburg-Essen, Essen, NRW, Germany; Department of Biomedical Informatics, University of Utah, Salt Lake City, UT, USA

## Abstract

Genomics research is complicated by the inherent difficulty of collecting, transforming, and integrating the numerous datasets and annotations germane to one’s research. Furthermore, these data exist in disparate sources, and are stored in numerous, often abused formats from multiple genome builds. Since these complexities waste time, inhibit reproducibility, and curtail research creativity, we developed Go Get Data (GGD; https://gogetdata.github.io/) as a fast, reproducible approach to installing standardized data recipes.

## Main

There is a need to standardize and simplify access to genomic data to enable reproducibility, remove common barriers to research, and foster studies that integrate diverse datasets. We developed Go Get Data (GGD) to address these challenges. Our approach is inspired by software package managers (e.g., pip (https://pip.pypa.io/en/stable/), Conda (https://conda.io), HomeBrew), which are popular because they use recipes to simplify and automate software installation via standard naming, version tracking, and dependency handling. We realized that the concept of a recipe could also be used to automatically locate, transform, standardize, and install *datasets.* GGD builds upon the software package framework in Conda, while modifications within GGD allow the Conda infrastructure to support datasets in addition to software. We chose to use Conda because of its wide acceptance and popularity within the life sciences with the support of Bioconda^1^, its version tracking and dependency handling, and its ability to normalize the installation of software packages across operating systems. Furthermore, Conda removes the dependence of administrative software management by installing all the desired packages within an isolated environment on a user’s system.

Conda provides a mature framework on which to build; however, managing genomic data comes with a unique set of challenges not seen with software management. GGD’s data recipes require additional knowledge regarding the version and provenance of the datasets, along with details about what makes the recipe unique. For example, genomics data is plagued by many inconsistencies such as genome build, chromosome labeling, sorting, indexing, and more, all of which require consistency and standardization in order to be properly managed. The resulting file format (e.g., BAM^2^, VCF^3^, BED^4^) must be correct, verified, and standardized for interoperability with common software and other datasets and annotations. Data processing is commonly required in order to use a dataset for an analysis. Processing genomic data also typically requires additional bioinformatics software tools, supplementary genomic datasets, and multiple curation steps. Storing the data recipes, associated metadata, genomic data files, and more all require a framework for access and management. In order to facilitate the use of data after installation, data files must be consistently organized and have a unique environment variable that points to the specific data package. These difficulties complicate genomic data management and are accounted for within the GGD framework.

Each GGD data recipe is a modified Conda recipe. Conda’s recipe format entails a powerful and sometimes complex set of possible statements, only a subset of which is relevant for data recipes. Therefore, in order to simplify the process, GGD partially automates the creation of a dataset recipe. Rather than requiring researchers to create all of the pieces required for the recipe, GGD only requires one to supply a bash script (**Figure 1A**). The bash script must contain the necessary steps for obtaining and transforming the raw data into a standardized data recipe. Once a bash script is provided, GGD command-line tools create (via ‘`ggd make-recipe`’) and validate (via ‘`ggd check-recipe`’) the recipe for use within the GGD and Conda frameworks.

**Figure 1.**
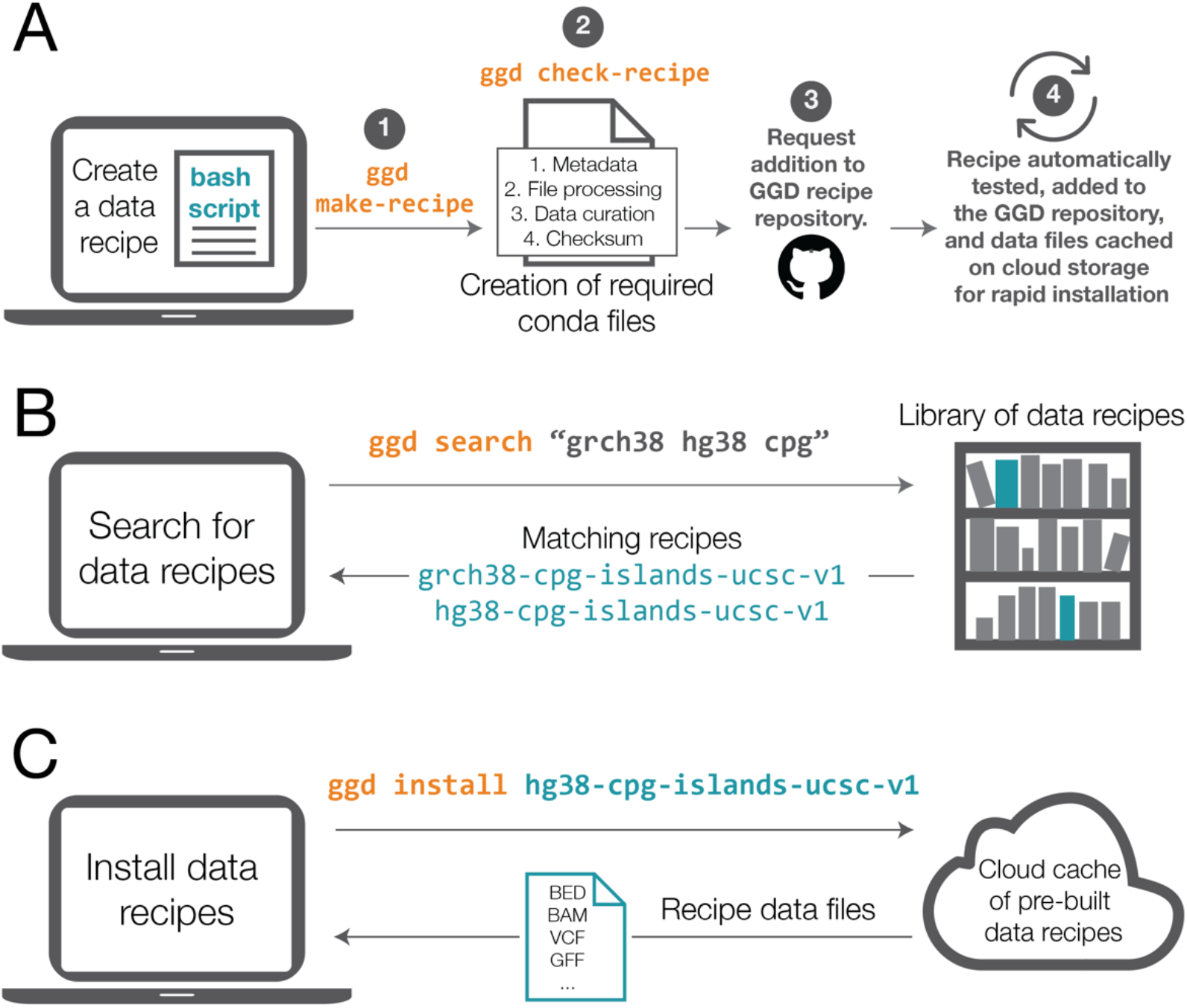
Overview of the creation and use of GGD data recipes. **A.** GGD creates a data recipe from a bash script, which defines the steps taken to access, process, and curate the desired data files. (1) The “ggd make-recipe” command incorporates the bash script and additional auto-generated files into a complete data recipe. (2) The “ggd check-recipe” command executes required tests and validates the created data recipe. (3) Once a data recipe has been tested and validated it can be added to the GGD data recipe repository on GitHub. (4) Each data recipe is further tested via an automatic continuous integration system. If validated, the recipe is transitioned into a data package, which is added to the Anaconda Cloud and the resulting data files are cached on AWS storage. **B.** Validated data packages can be found via the GGD command-line interface. For example, to find all data packages associated with “grch38” or “hg38” and the keyword “cpg” one would use “ggd search” with “grch38”, “hg38”, and “cpg” as search terms. GGD will identify and return all data packages within the GGD library that are associated with the search terms provided. **C.** The desired data package is installed via the “ggd install” command. If the data files are cached, they are downloaded directly. If the data package must be built from the recipe, GGD follows the instructions within the recipe while accounting for both software and data dependencies. Installation ends with tracking the version of the installed data package and the creation of local environment variables that facilitate the use of installed data packages.

A GGD recipe contains the information required to find and install the dataset, and to manage the resulting data recipe on a user’s system. Each GGD recipe contains a metadata file, a system processing script, a data curation script, and a checksum file (**Figure 1A**). The metadata file describes data package information such as software and data dependencies. It also tracks essential attributes such as the species, genome build, data provider, data version, and genomic file type. The system processing script provides recipe, metadata, and local file handling within the Conda environment, initiates data curation, and adds local environment variables for easy data file access. The data curation script provides the necessary steps to access, download, process, and install the data recipe. Finally, the checksum file is used to verify that the data files along with their content are installed as expected. Collectively, these files represent the instruction manual that enables GGD to automatically find, transform, install, and manage genomic data within a local Conda environment on a user’s system.

Once a GGD recipe has been created and tested, the recipe is added to the GGD data recipe repository on GitHub. A continuous integration system is used to automatically test and package recipes. Once packaged, the continuous integration system caches the resulting data files on cloud storage (currently AWS S3) for rapid user installation, uploads the packaged data recipe as a data package to the Anaconda cloud, and creates the metadata files necessary for use with GGD. This continuous integration system ensures the validity of each recipe and provides an automated approach that simplifies manual review of each data recipe added to GGD.

To install a data package, a researcher using GGD searches for a dataset or annotation by name and/or keyword (**Figure 1B**). Once the relevant data package is identified, the researcher uses GGD to install it and integrate it into their research (**Figure 1C**). One widely known disadvantage of Conda is the time required to identify all of the dependencies that a software package needs prior to installation (which has, however, been significantly improved in recent releases). Therefore, GGD has optimized the process of installing precomputed data recipes by bypassing Conda’s environment solving step. Instead, GGD directly installs prevalidated data recipes that have been cached on cloud storage, allowing fast data recipe installation (typically in 10 seconds or less).

Rapid, standardized installation of datasets and annotations removes many common frustrations that researchers face for common analyses. In turn, this increased simplicity allows one to quickly leverage multiple data packages and combine them with analysis software to address research questions (**Figure 2**). Furthermore, GGD can easily be integrated into reproducible processing and analysis workflow systems such as Nextflow^5^ and Snakemake^6^.

**Figure 2.**
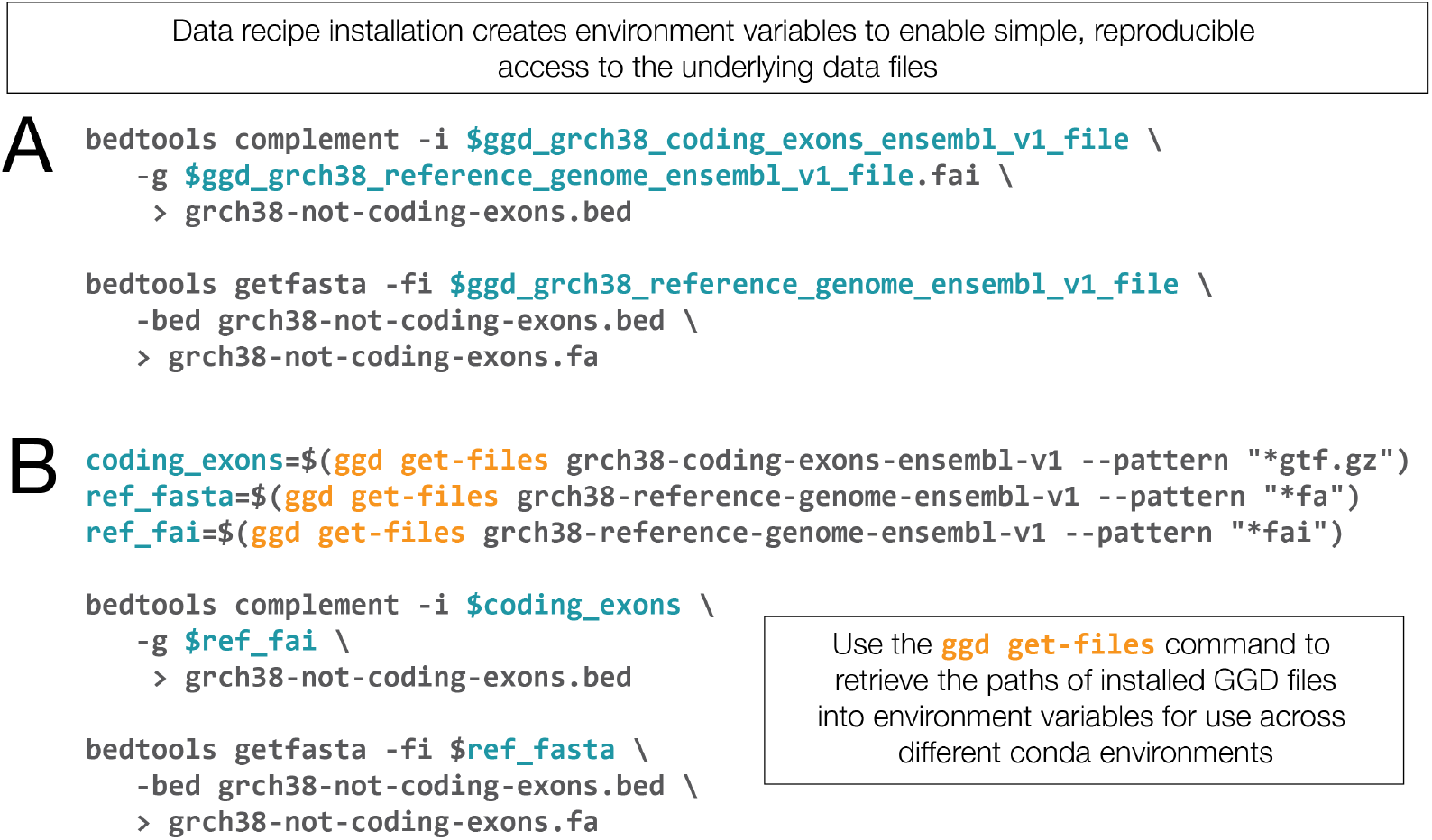
Using GGD data packages. **A.** Data recipe environment variables allow one to use the installed data files without needing to know where the files are stored or how to get them. For example, if one installed the `grch38-coding-exons-ensembl-v1` and `grch38-reference-genome-ensembl-v1` data packages, one could identify the complement between coding exons and a reference genome using each data file’s unique environment variable with the “`bedtools complement`” command. These environment variables allow one to perform any number of analyses with different bioinformatic tools or scripts. **B.** Using the “`get-files`” command, one can perform the same analysis on coding exons as seen in panel A. With data package environment variables, one needs to be in the environment where the packages were installed in order to use them. Alternatively, the “`get-files`” command provides access to data files installed by GGD and stored in either the currently active conda environment or a different non-active conda environment. Accessing data files in different environments is supported by the “--prefix” argument. This allows a user to install and store all data packages in a single conda environment while being able to access them from any other environment where GGD is installed.

Information about where the data originated, the version of the data being used, how it was processed, and distinguishing metadata (e.g., genome build for genomic recipes) are key components for every data recipe. This information acts as a unique identifier for the data recipe, and ensures data provenance and reproducibility. GGD maintains this information for each data recipe and provides multiple ways to obtain it through the documentation page for the recipe, the recipe stored in the ggd-recipes repo, or the (‘`ggd pkg-info`’) command (**Table 1**).

**Table 1.** A catalog of GGD tools available via the command-line interface.

| GGD CLI tool | Description of functionality |
| --- | --- |
| `ggd search` | Search for available data packages based on a search term(s) with genomic specific filters |
| `ggd predict-path` | Predict the installed file path for a file in a data package that has not been installed. (Useful for workflows like Snakemake) |
| `ggd install` | Install one or more data package(s) to a specific Conda environment on a user's system |
| `ggd uninstall` | Uninstall a data package from a Conda environment from a user's system |
| `ggd list` | Report the installed data packages within a specific Conda environment (similar to `conda list`) |
| `ggd get-files` | List the file(s) associated with an installed package on a user's system |
| `ggd pkg-info` | Retrieve the data package information of an installed data package on a user's system |
| `ggd show-env` | Display the GGD specific environment variables for the installed packages in a specific Conda environment on a user's system. |
| `ggd make-recipe` | Make a ggd data recipe that can be added to the ggd recipe ecosystem. The tool will transform a simple bash script into a ggd data recipe. |
| `ggd check-recipe` | Transform a ggd recipe that has been created from running `make-recipe` into a ggd data package and test the validity of the data package. |

Large-scale data integration is essential in all areas of genome research, since new annotations, datasets, and file formats are constantly being released. Through a suite of command-line tools (**Table 1**), GGD provides a standardized system for quickly finding, installing, managing, and creating data recipes.

GGD is a natural solution for enabling programmatic access to data in both ad hoc analyses and in more involved, frequently-used workflows. Whether within a container or on a local system, GGD can be used to install data packages before, during, or after the workflow process starts. Environment variables specific to the installed GGD data packages or use of the GGD command line interface can be used to access the data files for the desired process within the workflow. While currently focused on genomics datasets, the GGD framework has the capacity to support data management across many scientific disciplines. We expect that GGD will help to establish a standard, community-driven ecosystem for reproducible access to genomic and other scientific data. We encourage contributions from researchers to provide a comprehensive collection of reproducible data recipes to the scientific community. Further information can be found in the GGD documentation available at https://gogetdata.github.io/.

## References

1. Grüning B, Dale R, Sjödin A, Chapman BA, Rowe J, Tomkins-Tinch CH, Valieris R, Köster J, Bioconda Team. Bioconda: sustainable and comprehensive software distribution for the life sciences. Nat Methods. 2018 Jul;15(7):475–6. doi:10.1038/s41592-018-0046-7 PMID:29967506.

2. Li H, Handsaker B, Wysoker A, Fennell T, et al. The Sequence Alignment/Map format and SAMtools. Bioinformatics. 2009 Aug 15;25(16):2078–9. doi:10.1093/bioinformatics/btp352 PMID:19505943.

3. Danecek P, Auton A, Abecasis G, Albers CA, et al. The variant call format and VCFtools. Bioinformatics. 2011 Aug 1;27(15):2156–8. doi:10.1093/bioinformatics/btr330 PMID:21653522.

4. Kent WJ, Sugnet CW, Furey TS, Roskin KM, Pringle TH, Zahler AM, Haussler D. The human genome browser at UCSC. Genome Res. 2002 Jun;12(6):996–1006. doi:10.1101/gr.229102 PMID:12045153.

5. Di Tommaso P, Chatzou M, Floden EW, Barja PP, Palumbo E, Notredame C. Nextflow enables reproducible computational workflows. Nat Biotechnol. 2017;35(4):316–9. doi:10.1038/nbt.3820 PMID:28398311.

6. Koster J, Rahmann S. Snakemake--a scalable bioinformatics workflow engine. Bioinformatics. 2012 Oct 1;28(19):2520–2. doi:10.1093/bioinformatics/bts480 PMID:22908215.

